# Explainable autoencoder-based representation learning for gene expression data

**DOI:** 10.1101/2021.12.21.473742

**Authors:** Yang Yu, Pathum Kossinna, Qing Li, Wenyuan Liao, Qingrun Zhang

**Affiliations:** Department of Mathematics & Statistics, University of Calgary, Calgary, AB, T2N1N4, Canada; Department of Biochemistry & Molecular Biology, University of Calgary, Calgary, AB, T2N1N4, Canada; Alberta Children’s Hospital Research Institute, University of Calgary, Calgary, AB, T2N1N4, Canada

## Abstract

Modern machine learning methods have been extensively utilized in gene expression data analysis. In particular, autoencoders (AE) have been employed in processing noisy and heterogenous RNA-Seq data. However, AEs usually lead to “black-box” hidden variables difficult to interpret, hindering downstream experimental validation and clinical translation. To bridge the gap between complicated models and biological interpretations, we developed a tool, XAE4Exp (e**X**plainable **A**uto**E**ncoder for **Exp**ression data), which integrates AE and SHapley Additive exPlanations (SHAP), a flagship technique in the field of eXplainable AI (XAI). It quantitatively evaluates the contributions of each gene to the hidden structure learned by an AE, substantially improving the expandability of AE outcomes. By applying XAE4Exp to The Cancer Genome Atlas (TCGA) breast cancer gene expression data, we identified genes that are not differentially expressed, and pathways in various cancer-related classes. This tool will enable researchers and practitioners to analyze high-dimensional expression data intuitively, paving the way towards broader uses of deep learning.

**Availability:** Open source at https://github.com/QingrunZhangLab/Explainable-Deep-Autoencoder.

## Introduction

Genomic data, especially gene expressions, contain noise from known and unknown sources. As a result, many computational tools have been developed for preprocessing based on statistical methods such as linear mixed models and principal component analysis (Stegle et al., 2012). Recently, with the emerging success of modern machine learning, many researchers started to employ deep learning models for expression data processing (Koumakis et al., 2020). Due to its ability to deconvolute nonlinear factors and its unsupervised nature (Goodfellow et al., 2016), *autoencoders* (AE) have been extensively utilized to develop various tools for processing expression data (Taroni et al., 2017; Dwivedi et al., 2020).

A notable feature of AEs is their ability to learn the hidden *representations* of input data despite of the input being noisy and heterogenous, leading to “hidden variables” (**Supp. Fig. S1**) that are cleaner and more orthogonal for downstream analysis. However, such learned representations, although enjoying desirable statistical properties, are difficult to interpret. Therefore, in practice, researchers and clinical practitioners are left with manual inspection of data to decide whether to conduct costly experimental follow-up or clinical investigations.

As such, to learn representations of expression data, interpretable tools are urgently needed to accommodate explainable analyses. Following the efforts in machine learning, bioinformaticians have developed a couple of tools towards this line. In particular, Hanczar and colleagues employed gradient methods to analyze the contribution of specific neurons in the network (Hanczar et al., 2020), which however, does not focus on the contribution of inputs. Dwivedi and colleagues analyzed the effect of an input feature by differentiating the outcome by switching off the focal input feature, i.e., gene (Dwivedi et al., 2020). Although aiming to relieve the problem of interpretation, this method only focuses on the marginal effect of each gene, which does not employ the latest development in an emerging field, eXplainable Artificial Intelligence (XAI) that systematically examines complex models.

In XAI, SHapley Additive exPlanations (SHAP) (Lundberg et al., 2017) is a pioneering method inspired by the popular economic concept of “Shapley Value”, quantifying the contribution of a player in a game. (Awarded Nobel Prize in Economics in 2012). In brief, SHAP learns a linear model to approximate a complicated neural network to quantify the contribution of each input feature (**Supp. Fig. S2**). SHAP has been extended to many variants (Lundberg et al., 2020) and broadly used.

In this work, we developed a tool, XAE4Exp (e**X**plainable **A**uto**E**ncoder for **Exp**ression data) to facilitate a SHAP-supported explainable AE-based representation learning. Moreover, by applying it to real data, we reveal sensible pathways through SHAP-based explanations.

## Methods

The descriptions of AE, SHAP and XAE4Exp are detailed in **Supplementary Notes**. Here we briefly outline the implementation of XAE4Exp.

After initial quality control (QC), XAE4Exp conducts further QC by filtering genes with low variance. A DeepAE (Lore et al., 2017) with three hidden layers is used for representation learning (**Supp. Fig. S3(A)**). Then the Tree Explainer implementation of SHAP based on Random Forest (Lundberg et al., 2020) has been directed to the AE output to quantify the contribution of each gene to each hidden variable (i.e., representations) (**Supp. Fig. S3(B)**). Finally, for genes contributing substantially to a hidden variable (specified by a user-defined parameter, or defaulted as variables with nonzero SHAP values), XAE4Exp employs Over-Representation Analysis to conduct enrichment analysis (Subramanian et al., 2005) to characterize underlying pathways contributed by the disclosed genes (**Supp. Fig. S3(C)**).

The number of hidden variables in each hidden layer and optimizer selection of DeepAE are tuned based on the mean square error loss function (**Supp. Notes**). In addition, the number of hidden variables in the smallest hidden layer is selected based on the downstream pathway enrichment results. Overall, different parameters didn’t affect the results significantly.

To demonstrate the use of XAE4Exp, we applied it to The Cancer Genome Atlas (TCGA) breast cancer tumor tissue expression data (sample size N= 1041, number of genes M= 56,497).

## Results

In this demonstration (**Supp. Notes**), we chose the number of hidden variables in the smallest hidden layer of the AE to be 6. After learning, the R^2^ between input data and recovered data is 97.67%, sufficiently high to indicate that the training process went well.

The contributions of individual genes towards each hidden variable are revealed by SHAP and examples for a hidden layer is depicted in **Supp. Figs. S4 & S5**. Notably, most of SHAP-listed top genes are not differentially expressed (**Supp. Table S1**), indicating their roles in capturing the nonlinear contribution revealed by AE. For example, CUEDC2 has been experimentally verified that it interacts with ER-α, confers endocrine resistance in breast cancer (Pan et al., 2011). TRIM28 is associated with Wilms Cancer 1 and 5, and related to DNA Damage and Corticotropin-releasing hormone signaling pathways etc.

By conducting enrichment analysis using genes with their nonzero SHAP values, we identified 126 enriched pathways (**Supp. Table S2**). As comparisons, we also carried out differential expression or DE (Ritchie et al., 2015) and differential co-expression, or DiffCoEx (Tesson et al., 2010) analysis (**Supp. Notes**). The outcomes are presented in (**Supp. Table S3 & S4**). Overall, XAE4Exp shared around 30% and 52% with DE and DiffCoEx, respectively. However, AE4Exp picked substantially more (84) pathways (**Supp. Fig. S6**), especially in the classes of “Cancer: specific types”, “Cancer: overview”, “Cell Growth and Death”, “Infectious Disease” and “Immune System” (**Supp. Table S5**). XAE4Exp’s outcome has larger overlap with network-based approach DiffCoEx, possibly because that AE can handle nonlinear relationships among features, whereas traditional DE analysis couldn’t utilize gene network information.

## Conclusion and Discussion

Although deep neural networks are famous for being sample hungry, our study showed that DeepAE could be used for “small sample-sized” expression data, if supported by XAI to quantify the contribution of input features. Importantly, genes not differentially expressed were also revealed by our tool, taking advantage of AE’s ability to handle nonlinear structure. In summary, XAE4Exp is a useful tool for researchers to interpret the AE-based data preprocessing, leading to insightful knowledge on how to utilize the representations revealed by the AE-based machine learning. In this work, only DeepAE is used. However, the tool could be straightforwardly extended to other AEs such as Variational AE for causal inference, and Graph AE to learn causal structure, which is our planned future work. Similarly, in addition to the Random Forest explainer, other explainers such as Gradient Boosting Decision Tree (GBDT), CatBoost, etc can also be incorporated straightforwardly.

## Acknowledgement

Supported by an NSERC Discovery Grant (RGPIN-2018-05147), an NSERC CRD grant (CRDPJ532227-18) and NFRF Exploration grant (NFRFE-2018-00748).

## Supplementary Materials

### Data source and specifications of computing infrastructure

#### Data source

For benchmarking on real data, we used a public dataset: The Cancer Genome Atlas (TCGA) breast cancer tumor tissue expression data (sample size N= 1041, number of genes M= 56497). The raw count data was downloaded from the TCGA data portal and then converted to Transcripts Per Million (TPM) values using gene lengths obtained through the BioMart package. We then performed basic quality control to examine the overall structure of the data by PCA analysis and did not find unexpected data points.

The following steps were taken to obtain the data source:

- Go to https://portal.gdc.cancer.gov/
- Click on the Exploration tab
- Check the “TCGA-BRCA” and “ductal and lobular neoplasms” options on the left.
- Click “View files in Repository”
- Under “Workflow Type” in the left, select “HTSeq-Counts”.
- Click the “Add All Files to Cart” button on the right.
- Then, go to “Cart” on the upper right-hand side of the top bar.
- Download the sample sheet (this indicates the type of sample) and then click Download: Cart.

Similar process for other cancers as well. Once downloaded, the compressed files need to be extracted and put in one folder (the ‘.count’ files). Then convert the count data to the TPM expression matrix (code is available in our GitHub). Please note that this still contains both tumor and normal samples. In this project, we only used tumor data.

#### Data processing

XAE4Exp conducts an initial quality control (QC) by filtering genes with variance lower than 1.0, which reduces interference from unimportant genes. After the initial quality control, the number of genes decreased from M= 56497 to M’ = 22920.

#### Specification of the computing server

GPU: NVIDIA Tesla V100 DGXS 32 GB @ 1297 MHz

Memory: 32 GB HBM2

Network: 897.0 GB/s

### Detailed algorithm for Deep Autoencoder

Autoencoder (AE) is an artificial neural network that learns to efficiently encode representations for unlabeled data and then using the learned representations to reconstruct back the original input as close as possible. It reduces the dimensionality by training the network to extract useful information and ignore noise (Kramer, 1991). The basic AE model is a simple three-layer neural network structure: an input layer, a hidden layer, and an output layer **(See Supplementary Figures S1)**. AE is a data compression algorithm in which the data compression and decompression functions are data-dependent, lossy, and automatically learned from samples.

#### AE consists of two parts

- Encoder: this part compresses the input into a potential spatial representation, which can be represented by the encoding function *h* = *f*(*x*)
- Decoder: this part reconstructs the input from the potential spatial representation and can be represented by the decoding function *r* = *g*(*h*)

Thus, the whole AE can be described by a function *g*(*f*(*x*)) = *r*, where the output *r* is similar to the original input *x*. The purpose of training AE is to get a useful *h* that can represent the input *x* well.

#### The ideal AE model takes into account the following points

- Sensitive enough to the input to accurately build a reconstruction.
- Insensitive enough to the input so that the model does not simply memorize or over-fit the training data.

This trade off forces the model to keep only the data changes needed to reconstruct the input, without retaining redundancy in the input. In most cases, this involves constructing a loss function. The loss function defines as *L*(*x, g*(*f*(*x*))), where *L* is the loss function to calculate the difference of *x* and *g*(*f*(*x*)). XAE4Exp uses Mean Square Error (MSE) as the loss function.

Deep Autoencoder (DeepAE) (Vaibhaw, 2018) is a variant of AE, it is composed of two symmetrical networks, usually with multiple shallow layers representing the encoded half and a second set of multiple layers forming the decoded half. Given enough data, DeepAE has the advantage of being able to both create relevant features from raw data and identify highly complex nonlinear relationships.

Compared to other machine learning models such as supervised learning models, DeepAE has the advantage of not requiring a lot of manual labelling. Meanwhile, DeepAE, designed in a deep neural network, has a good performance in both linear and nonlinear relationships, because DeepAE has the advantage of being able to create relevant features from raw data and identify highly complex nonlinear relationships if there are enough data. Gene expression has a complex regulation network, so that DeepAE could be an excellent match for gene expression data. (Sanjiv K. Dwivedi et al 2020). DeepAE has a better performance in nonlinear problems than AE, and it is more stable and faster than other variations of AE. That’s why DeepAE could be the best choice for the first study in this direction.

#### Mean Square Error (MSE) loss function

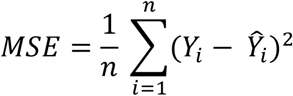

Where *n* is the number of data points, *Y_i_* represents observed input values, and 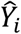 represents predicted values.

### Detailed algorithm of SHapley Additive explanation (SHAP)

SHapley Additive exPlanation (SHAP), as a classical post-hoc explanatory framework, can calculate the contribution value for each feature variable in each sample to achieve the explanatory effect (Lundberg and Lee, 2016). The value is specifically referred to as SHAP Value. Our existing dataset contains many feature variables, and from the perspective of game theory, each feature variable can be treated as a player. The prediction results obtained by training the model with this dataset can be regarded as the benefit of cooperation among many players to complete a project, see **Supplementary Figure S2** for an illustration.

When we perform a SHAP post-hoc interpretation of the model, we need to label it explicitly. The known dataset (with *M* feature variables and *N* samples), the original model *f*, and all predicted values of the original model *f* on the dataset. *g* is the model used to interpret *f* in SHAP. The dataset is firstly predicted using *f* to obtain the mean of the model predicted values *ϕ*_0_. A single sample is denoted as *x* = (*x*_1_, *x*_2_, *x*_3_ … *x_m_*) and *f*(*x*) is the predicted value under the original model. *g*(*x*) is the predicted value of the post hoc explanatory model, satisfying

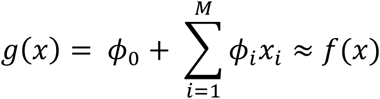

where *ϕ_i_* represents the SHAP Value of the *ith* feature variable, which is the core value to be calculated in SHAP and needs to satisfy uniqueness (Lundberg, et al., 2020).

#### The above model g enjoys the following properties

- **Local accuracy:** The predicted values obtained by the two models are equal. When a single sample *x* is input to model *g*, the predicted value 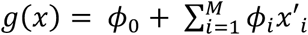 obtained is approximately equal to the predicted value *f*(*x*) obtained by the original model.
- **Missingness:** If there is a missing value in a single sample, i.e., no value is taken under a characteristic variable that has no effect on the model g, its SHAP Value is 0.
- **Consistency**: If a change in a model increases or keeps constant the contribution of certain features without considering other inputs, the attribution of that input should not be reduced.

Under the above three constraints, it can be proved theoretically that the following expressions *ϕ_i_*, as well as the corresponding model *g*, are unique:

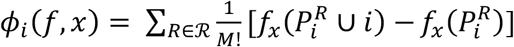

where 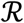 is the set of all feature orderings, 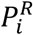 is the set of all features that come before feature *i* in ordering *R*, and *M* is the number of input features for the model (Lundberg, et al., 2020).

#### TreeSHAP

TreeSHAP is a SHAP method specifically for the tree model. Tree integration models include many black box models with good performance, such as Random Forest, XGBoost, LightGBM and CatBoost, all of which are nonlinear models. XAE4Exp uses Random Forest as the original model *f* and Tree Explainer as the model *g* to interpret *f* in SHAP.

#### Random Forest

Random Forest is an extension of the bagging algorithm in Ensemble Learning. A random forest is composed of many decision trees, and different decision trees are not associated with each other. When we perform the classification task, new input samples enter and let each decision tree in the forest judge and classify them separately. Each decision tree will provide a classification result of its own. Whichever classification result of the decision tree has the most supports, then the random forest will take this classification as the final output. This allows efficient training of the model when the sample feature dimension is high, which works very well for high dimensional transcriptome data.

### Parameters and commands for different tools

We used the default parameters of XAE4Exp. According to DeepAE (Vaibhaw, 2018) and SHAP (Dataman, 2019), specified hidden layer of DeepAE and SHAP TreeExplainer with Random Forest Regressor were used for real data analysis.

#### DeepAE (with Quality Control) parameters and explanation

~~~
*#Model name*
model_name = ‘AE_Geno’
*#Absolute path for breast cancer data txt file*
PATH_TO_DATA = ‘./data.txt’
*#Absolute path to save representation result from DeepAE*
PATH_TO_SAVE = ‘./AE.txt’.
*#Name of data type*
RNA_name = ‘BRCA’
*#Quality control by filtering genes which variance less than 1*
geno_var[i]<1
*#Number of samples selected for one training session*
batch_size = 4096
*#Test sample selection ratio, which also decides the train_size = 1 – test_size*
test_size=0.1
*#Number of methods for generating random numbers drawn from training, testing data*
random_state=42
*#Number of processes run at the same time (the maximum number depends on GPU/CPU*)
num_workers=8
*#Gene number of cleaned data*
snp = int(len(geno[0]))
*#Number of hidden variables (representation*)
smallest_layer = int(snp / 20000)
*#Number of hidden nodes of First and Third hidden layer*
hidden_layer = int(2*smallest_layer)
*#Loss function type*
distance = nn.MSELoss()
*#Optimizer type*
optimizer = torch.optim.Adam()
*#Learn rate of optimizer*
lr=0.0001
*#Round number of training*
num_epochs = 200
*#Initial loss in training processes*
sum_loss = 0
*#Initial batch number in training processes*
current_batch = 0
*#Initial loss in testing processes*
test_sum_loss = 0
*#Initial batch number in testing processes*
test_current_batch = 0
~~~

#### Random Forest Regressor and SHAP figures/txt parameters and explanation

~~~
*#Absolute path for cleaned breast cancer data txt file*
PATH_TO_DATA = ‘./data_QC.txt’
*#Absolute path for DeepAE representation of the last epoch*
PATH_TO_AE_RESULT = ‘./AE_199.txt’
*#Absolute path to save SHAP figure*
PATH_TO_SAVE_FIGURE = ‘.figure.pdf’
*#Absolute path to save SHAP txt*
PATH_TO_SAVE = ‘./shap.txt’.
*#Initial summary SHAP value*
total_value = 0
*# Test sample selection ratio, which also decides the train_size = 1 – test_size*
test_size=0.2
*#Number of methods for generating random numbers drawn from training, testing data*
random_state=42
*#Depth of each tree in the forest*
max_depth=20
*#Absolute path for breast cancer data txt file*
random_state=42
*#Number of trees in the forest*
n_estimators=100
*#Initial chart for SHAP value, which shows impact of every sample with every feature*
shap.summary_plot(shap_values, X_test)
*#Bar chart for SHAP value, which represents mean /SHAP value/ for each gene*
shap.summary_plot(shap_values, X_test, plot_type=“bar”)
*# Name of data type*
RNA_name = ‘BRCA’
*#Autoencoder compress number*
compress_num = ‘1’
~~~

### XAE4Exp Processes

#### Data preparation

To demonstrate the use of XAE4Exp, we applied our method to the tumor tissue expression data (sample size N= 1041, number of genes M= 56497) of breast cancer from the TCGA project. After initial basic quality control of the data, we removed genes with variance lower than 1.0 As a result, the total number of genes is reduced from 56,497 to 22,920. Quality control helps us decrease the noise of the original data set, which is an essential step for data preprocessing. To make the further SHAP explainer clearer, we convert gene ids to HGNC symbols by an R package named biomaRt. It should be noted that there are a small number of gene ids that don’t have related **HGNC symbols**, those gene ids will be shown as ‘unnamed: gene id’ in the graphs.

#### DeepAE

A three-hidden-layer DeepAE, which uses mean square error (MSE) as the loss function and Adam as the optimizer, is used in XAE4Exp for representation learning, which compress the 22,920 feature variables (gene) to 6 hidden variables (representation). The validation accuracy of DeepAE reached 97.67%, which implies that the representation remains most information of original dataset.

It is important to note that for a specific representation, there are some hidden variables that are zero values because they represent noise in the original data.

#### Random Forest Regressor and SHAP explanation

XAE4Exp uses Random Forest as the original prediction model and TreeExplainer as the SHAP model to predict Random Forest results. The Random Forest Regressor builds on 100 decision trees (estimator) and each decision with a depth of 20. As a strong explainer, it generated SHAP value for every feature of every sample.

Based on non-zero representation learning result compressed by DeepAE, SHapley Additive exPlanations generated groups of feature importance figure (an example illustrated in **Supplementary Figures S4**), global interpretation figure (an example illustrated in **Supplementary Figures S5**) and SHAP values. The idea behind the importance of SHAP features is simple. Features with large absolute SHAP values are important. Since we wanted global importance, we added up the absolute SHAP values of each feature in the data, then we sorted the features (genes) by decreasing importance. Thus, we generated the gene module for further enrichment analysis by this feature importance of each gene.

#### Over-Representation Analysis

Over Representation Analysis (ORA) (Boyle et al. 2004) is widely used to determine whether known biological pathways or functions are enriched in a list of genes, for example, list of differentially expressed genes (DEGs). The p-value can be calculated by the hypergeometric distribution is

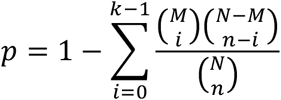

where N is the total number of genes in the background distribution, n represents the size of the list of genes of interest (for example the number of differentially expressed genes), M stands for the number of genes in the distribution (directly or indirectly) annotated as gene set of interest, and k is the number of genes in the list annotated to the gene set. The background distribution, by default, is all the genes that have annotation.

#### Results of XAE4Exp

The feature importance generated by SHapley Additive exPlanations method shows the critical features in decreasing order. Examples for a hidden layer is depicted in **Supp. Figures S4 and S5.** The CUEDC2 gene is ranked as the top important one, even though it is not differentially expressed. In Pan et al. 2011 paper published in Nature Medicine, they have experimentally identified that “CUEDC2 interacts with ER-α and promotes ER-α ubiquitination and degradation.” (Pan et al. 2011).

In this paper, genes with absolute Shapley value larger than zero were used for KEGG pathway enrichment analysis. XAE4Exp identified 126 pathways, shared with more than 52% of the pathways identified by differential co-expression analysis, and around 30% of the pathways picked by differential expression analysis (**Supplementary Figure S6**). Since autoencoder can handle both linear and non-linear transformations, it has higher power to capture gene cooperation systems. It is no surprise that XAE4Exp has an outcome close to that of networkbased approaches like differential co-expression. Differential expression can give us direct overview about which genes has significant change between two different conditions but is not informative to understand gene network.

Pathways identified by XAE4Exp are classified in a few major classes which are highly related to cancer pathogenesis, for example, “Cell growth and death”, “Specific cancer”, “Infectious disease” etc. **Supplementary Table S5** shows the number of pathways in each class. **Supplementary Table S2** lists the detailed 126 pathways. Pathways like “Apoptosis”, “Cell cycle”, “p53 signaling pathway”, “Cellular senescence”, “Ferroptosis”, “Base excision repair” etc. are highly related to cancer cell immortal status. Most of these pathways have been reported to be relevant to breast cancer pathogenesis.

Compared with DE and DiffCoEx, XAE4Exp identifies more pathways related to other cancers, infectious diseases, and immune systems etc. It brought us a broader view of the carcinogenesis.

### Comparison with Differential Expression (DE)

Single-gene-based differential expression (DE) analysis is the most popular method adapted by many researchers. To compare the performance of XAE4Exp to DE, we processed the same data set with Differential Expression. An edgeR-limma-based pipeline was used to normalize the data to log2-counts per million (log-CPM) values and a linear model incorporating weights to correct for the mean-variance relationship was used to statistically detect differential expression of genes. The pipeline was run using default values for all parameters as described in the workflow. We then performed gene set enrichment analysis followed by DE, and the results are shown in **Supplementary Table S1**. With loser FDR criteria (FDR <=0.1), we identified 28 pathways (shown in **Supplementary Table S3)**, 8 shared with XAE4Exp, 7 shared with DiffCoEx methods.

### Comparison with Differential Co-expression Analysis

To compare our method with a standard method based on network analysis, we used differential co-expression networks to identify groups (or “modules”) of differentially co-expressed genes and conduct pathway enrichment on these modules. While there are multiple methods of differential co-expression analysis, the most widely used method remains DiffCoEx (Tesson et all 2010), an extension of the popular WGCNA (Langfelder, 2008). DiffCoEx begins with the construction of two adjacency matrices: 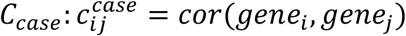 for cae samples and *C_control_* similarly for control samples.

While different correlation measures can be used in this step, the authors of DiffCoEx used the Spearman rank correlation. A matrix of adjacency difference is then calculated:

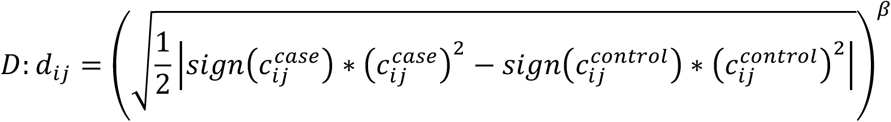

where *β* ≥ 0 is an integer tuning parameter which can be selected in multiple ways. In this paper we chose *β* ∈ [5,6,7,8,9,10] such that we can achieved minimum number of modules with the largest module containing the smallest number of genes. Next, a Topological Overlap dissimilarity Matrix is calculated where smaller values of *t_ij_* indicate that a pair of genes *gene_i_* and *gene_j_* have significant correlation changes (between case and control) with the same group of genes.

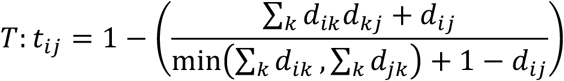

Finally, the dissimilarity matrix *T* is used for clustering and “modules” of differentially coexpressed genes are identified. These modules which contain sets of genes were then tested for pathway enrichment. **Supplementary Table S4** shows the ORA result of DiffCoEx.

### Supplementary Figures

**Supplementary Fig. S1:**
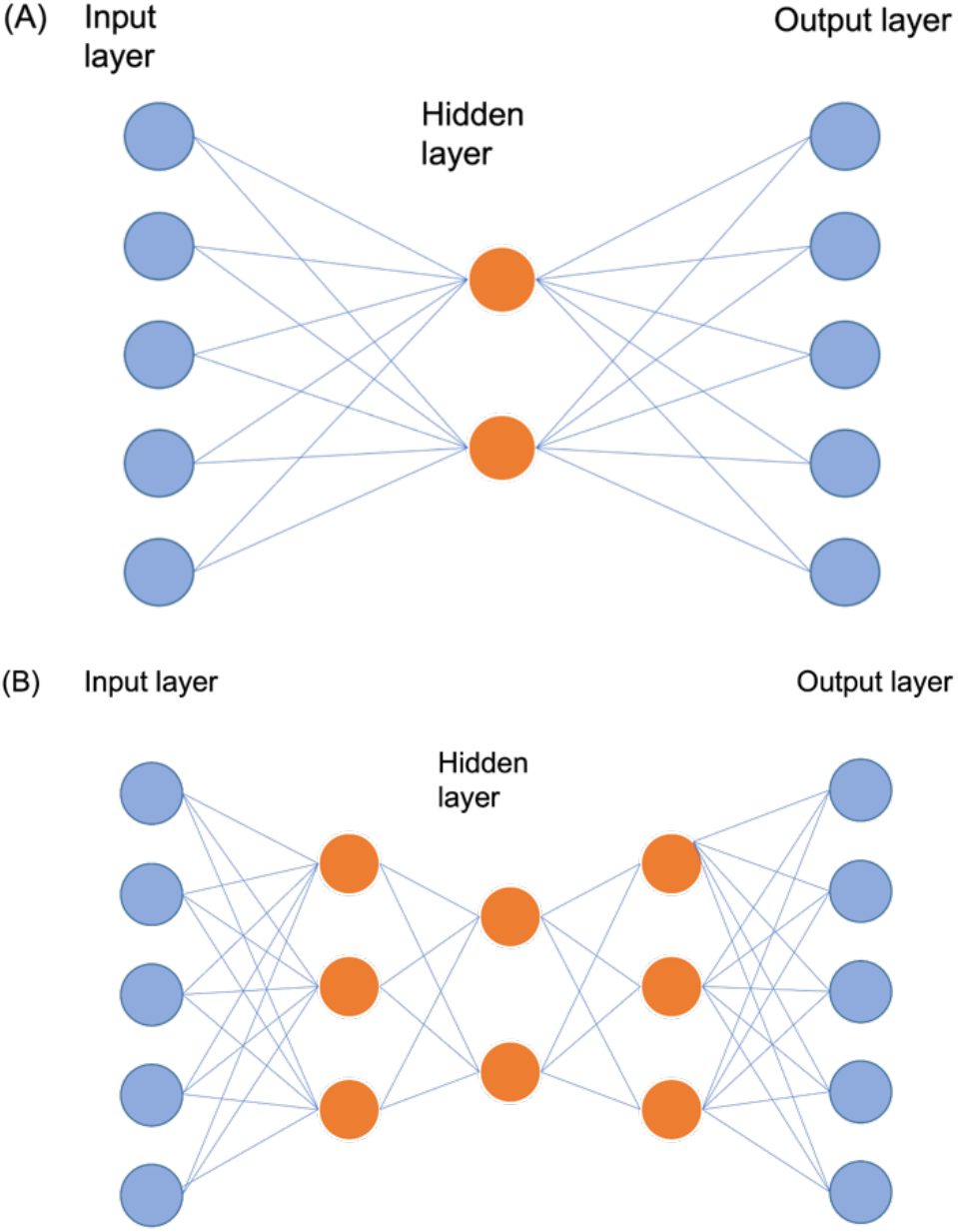
Autoencoder, or AE. **(A)** A simple AE with an input layer, a hidden layer, and an output layer. **(B)** A three hidden-layer Deep Autoencoder Neural Network. The output layer has the same number of dimensions as the input layer. The input and output of the AE are consistent, and the goal is to reconstruct itself using sparse higher-order feature recombination.

**Supplementary Fig. S2.**
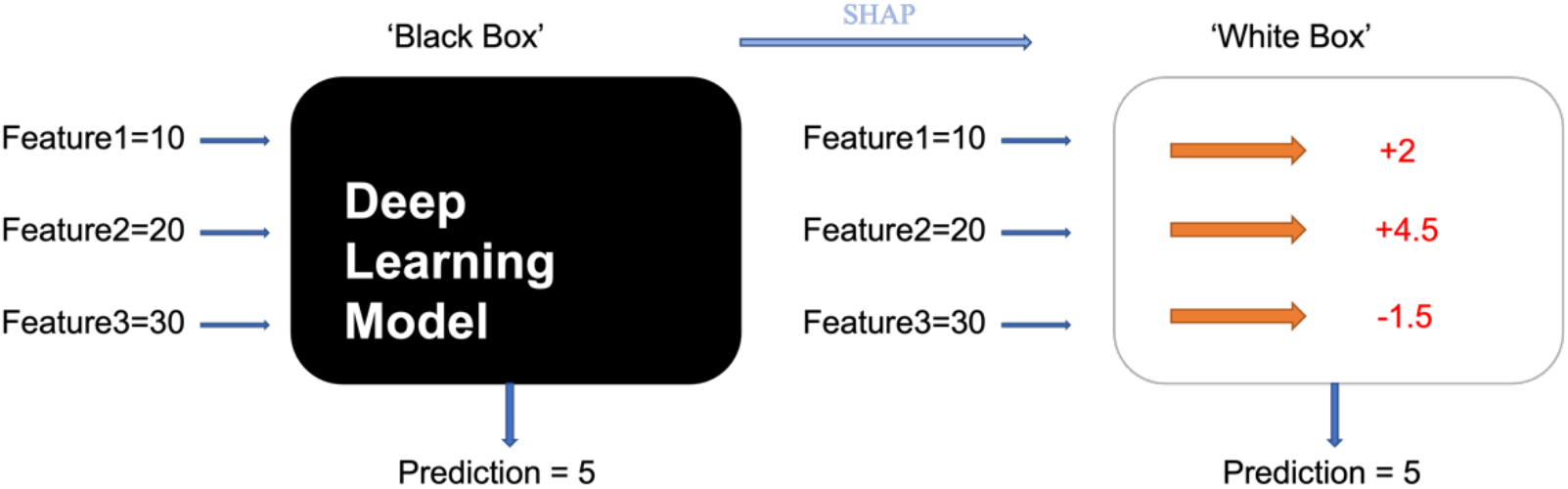
Illustration of SHAP framework. The black-box machine learning model has been explained by the quantified contribution of each input feature, by assigning a numeric measure of credit to each feature.

**Supplementary Fig. S3:**
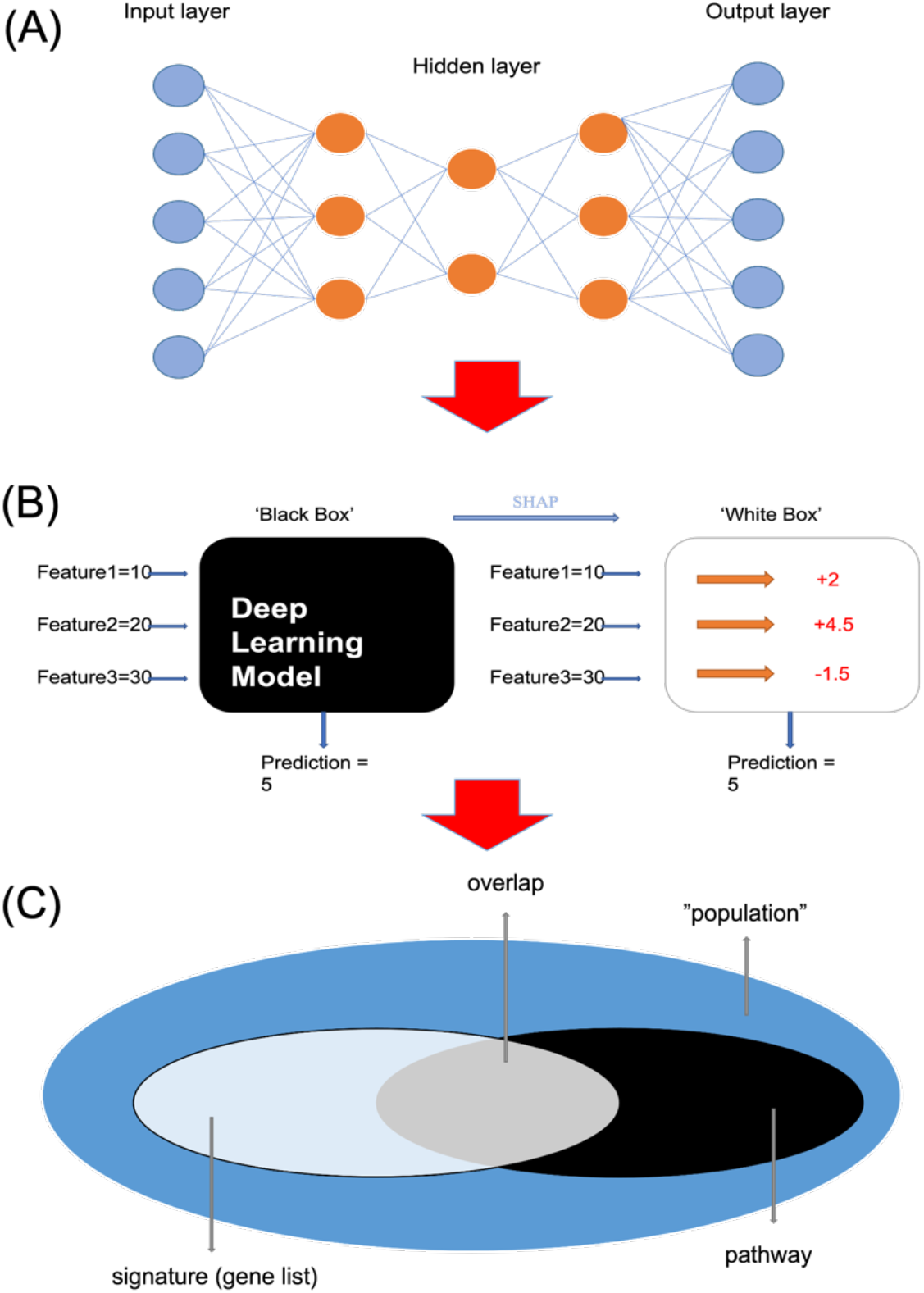
the XAE4Exp framework. **(A)** Step 1: A three hidden-layer DeepAE to train the representation. **(B)** Step 2: SHAP explanations enable a wide variety of new ways to understand global model structure. **(C)** Step 3: Enrichment analysis of a biologically relevant pathways of relevant genes.

**Supplementary Fig. S4.**
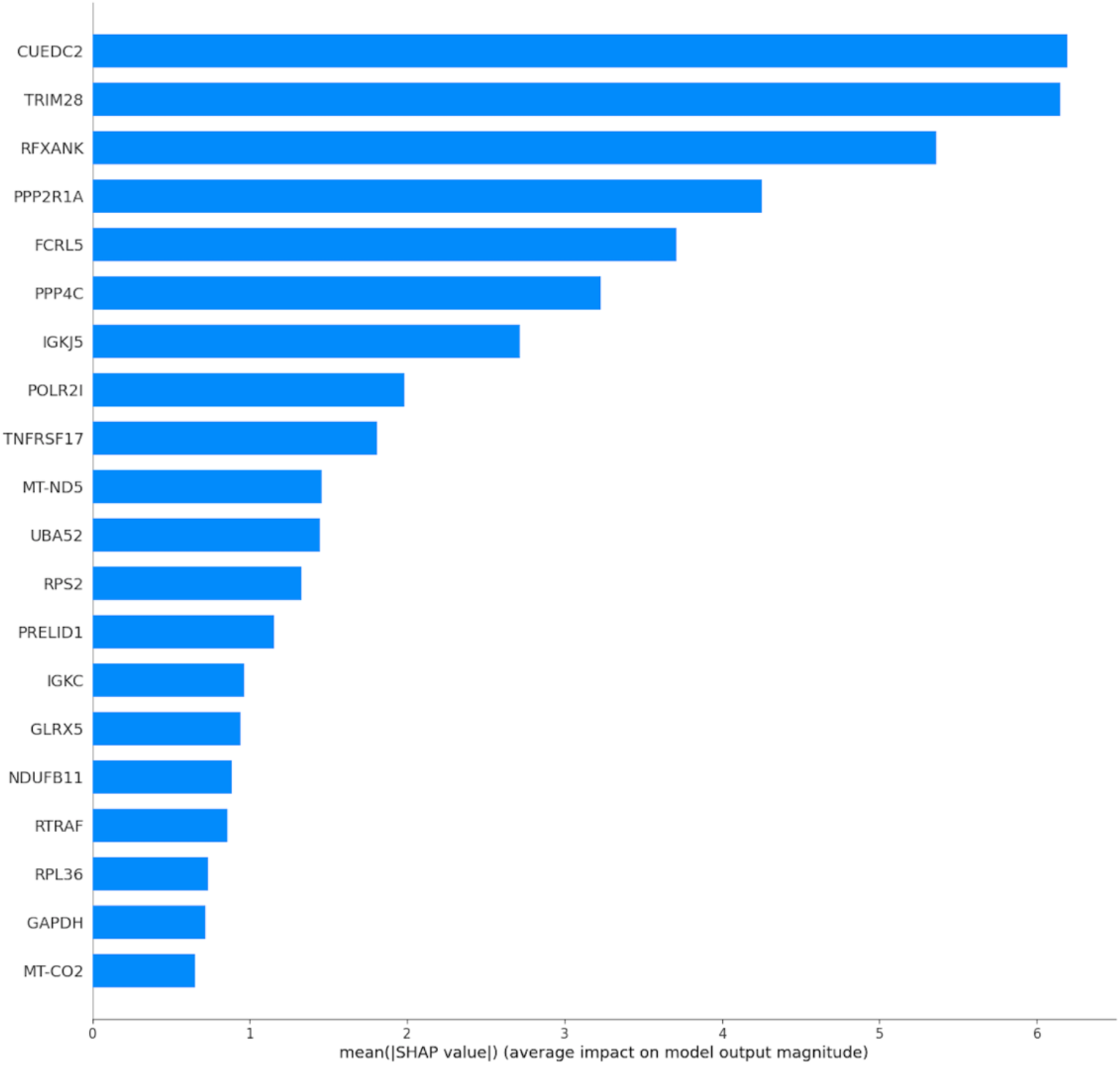
Top 20 features ranked by their importance assessed by SHAP value of a hidden variable among the total 6 hidden variables representations. Features with larger absolute SHAP values are more important. Since we wanted global importance, we added up the absolute SHAP values of each feature in the data, then features are sorted by decreasing order. CUEDC2 is ranked as the most important gene even though it is not differentially expressed. Researchers have experimentally identified that “CUEDC2 interacts with ER-α and promotes ER-α ubiquitination and degradation.” (Pan et al. 2011)

**Supplementary Fig. S5.**
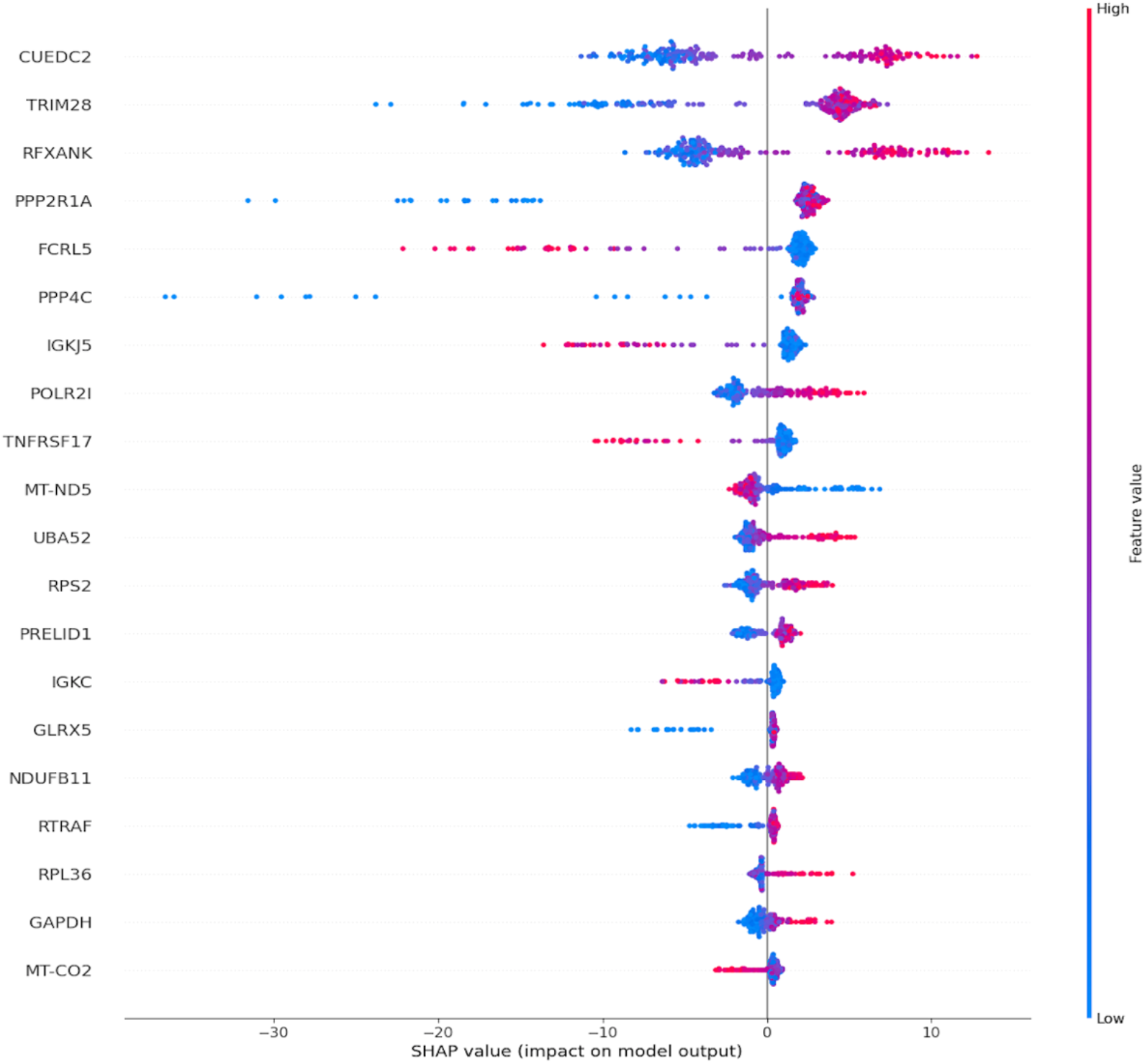
Top 20 global interpretation of SHAP value for a hidden variable. The feature determines the position on the y-axis, and the SHAP value determines the position on the x-axis. The colours represent the feature values from low to high (from blue to red). The vertical clustered points to reflect the distribution of SHAP values for each feature. These features are ranked according to their importance.

**Supplementary Fig. S6.**
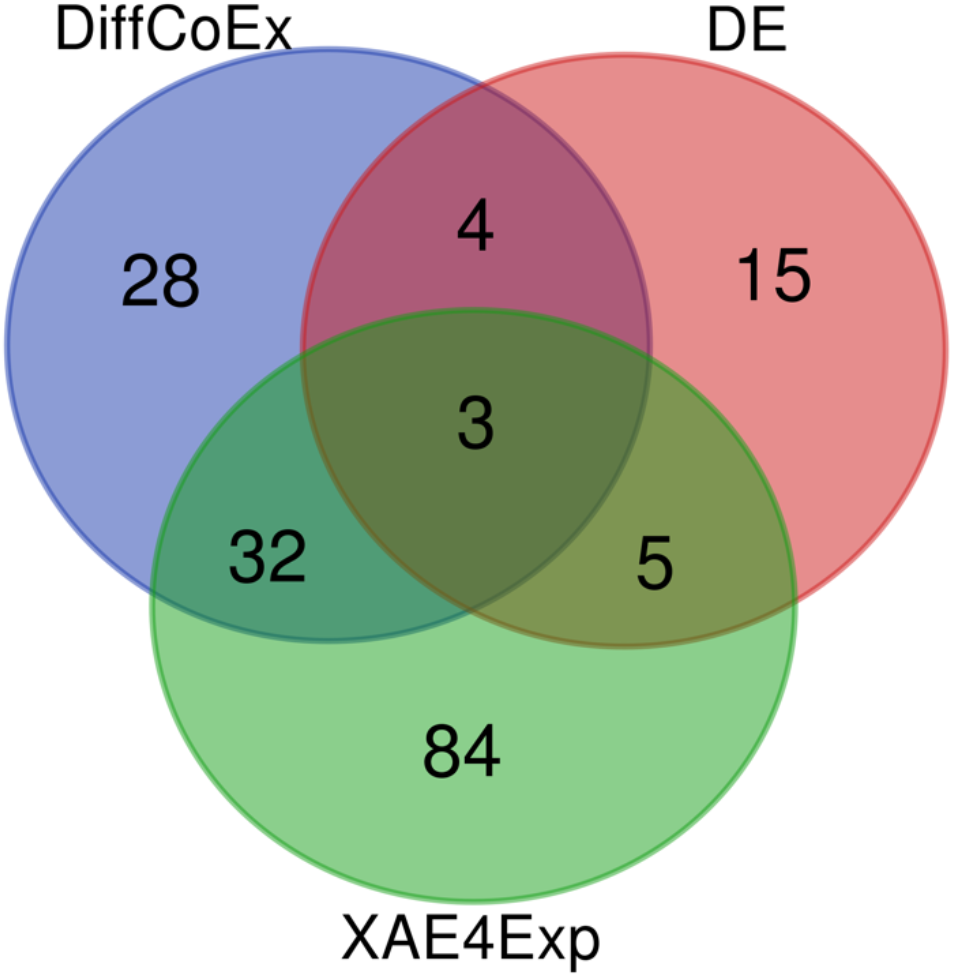
Comparison of XAE4Exp to Differential Expression (DE) and Differential Co-expression (DiffCoEx) by a Venn diagram.

### Supplementary Tables

Please refer to the Excel spreadsheet for Supplementary Tables S1-S5

## References

Dwivedi, S. K., Tjärnberg, A., Tegnér, J., & Gustafsson, M. (2020). Deriving disease modules from the compressed transcriptional space embedded in a deep autoencoder. Nature communications, 11(1), 1–10.

Goodfellow, I., Bengio, Y., & Courville, A. (2016). Deep learning. MIT press.

Hanczar, B., Zehraoui, F., Issa, T., & Arles, M. (2020). Biological interpretation of deep neural network for phenotype prediction based on gene expression. BMC bioinformatics, 21(1), 1–18.

Koumakis, L. (2020). Deep learning models in genomics; are we there yet?. Computational and Structural Biotechnology Journal.

Lore, K. G., Akintayo, A., & Sarkar, S. (2017). LLNet: A deep autoencoder approach to natural low-light image enhancement. Pattern Recognition, 61, 650–662.

Lundberg, S. M., & Lee, S. I. (2017). A unified approach to interpreting model predictions. In Proceedings of the 31st international conference on neural information processing systems (pp. 4768–4777).

Lundberg, S. M., Erion, G., Chen, H., DeGrave, A., Prutkin, J. M., Nair, B., … & Lee, S. I. (2020). From local explanations to global understanding with explainable AI for trees. Nature machine intelligence, 2(1), 56–67.

Meng, Q., Catchpoole, D., Skillicom, D., & Kennedy, P. J. (2017, May). Relational autoencoder for feature extraction. In 2017 International Joint Conference on Neural Networks (IJCNN) (pp. 364–371). IEEE.

Pan, X., Zhou, T., Tai, Y. H., Wang, C., Zhao, J., Cao, Y., … & Zhang, X. M. (2011). Elevated expression of CUEDC2 protein confers endocrine resistance in breast cancer. Nature medicine, 17(6), 708–714.

Ritchie, M. E., Phipson, B., Wu, D. I., Hu, Y., Law, C. W., Shi, W., & Smyth, G. K. (2015). limma powers differential expression analyses for RNA-sequencing and microarray studies. Nucleic acids research, 43(7), e47–e47.

Stegle, O., Parts, L., Piipari, M., Winn, J., & Durbin, R. (2012). Using probabilistic estimation of expression residuals (PEER) to obtain increased power and interpretability of gene expression analyses. Nature protocols, 7(3), 500.

Subramanian, A., Tamayo, P., Mootha, V. K., Mukherjee, S., Ebert, B. L., Gillette, M. A., … & Mesirov, J. P. (2005). Gene set enrichment analysis: a knowledge-based approach for interpreting genome-wide expression profiles. Proceedings of the National Academy of Sciences, 102(43), 15545–15550.

Taroni, J. N., Grayson, P. C., Hu, Q., Eddy, S., Kretzler, M., Merkel, P. A., & Greene, C. S. (2019). MultiPLIER: a transfer learning framework for transcriptomics reveals systemic features of rare disease. Cell systems, 8(5), 380–394.

Tesson, B. M., Breitling, R., & Jansen, R. C. (2010). DiffCoEx: a simple and sensitive method to find differentially coexpressed gene modules. BMC bioinformatics, 11(1), 1–9.

## References

Kramer, M. A. (1991). Nonlinear principal component analysis using autoassociative neural networks. AIChE journal, 37(2), 233–243.

Jordan, J. (2018). Introduction to autoencoders. Jeremy Jordan, Mar.

Lundberg, S., & Lee, S. I. (2016). An unexpected unity among methods for interpreting model predictions. arXiv preprint arXiv:1611.07478.

Vaibhaw, V. (2018). Building Autoencoder in Pytorch - Vipul Vaibhaw. Medium. https://vaibhaw-vipul.medium.com/building-autoencoder-in-pytorch-34052d1d280c

Dataman, D. (2019). Explain Any Models with the SHAP Values - Use the KernelExplainer. Medium. https://towardsdatascience.com/explain-any-models-with-the-shap-values-use-the-kernelexplainer-79de9464897a.

Boyle, E. I., Weng, S., Gollub, J., Jin, H., Botstein, D., Cherry, J. M., & Sherlock, G. (2004). GO:: TermFinder—open source software for accessing Gene Ontology information and finding significantly enriched Gene Ontology terms associated with a list of genes. Bioinformatics, 20(18), 3710–3715.

Langfelder, P., & Horvath, S. (2008). WGCNA: an R package for weighted correlation network analysis. BMC bioinformatics, 9(1), 1–13.

